# Bisphenol S causes deficits in social behaviour by disrupting serotonergic and BDNF-CREB1 signaling pathways

**DOI:** 10.64898/2026.06.20.733535

**Authors:** A K M Munzurul Hasan, Mahesh Rachamalla, Som Niyogi, Douglas P. Chivers

## Abstract

Bisphenol S (BPS), a widely used substitute for bisphenol A, is increasingly detected in aquatic environments; however, its neurodevelopmental effects remain insufficiently understood. This study investigated whether developmental exposure to an environmentally relevant concentration of BPS disrupts social behaviour and underlying neurobiological pathways in zebrafish *(Danio rerio).* At 21 days post-fertilization, BPS-exposed larvae exhibited a significant reduction in social preference, indicating impaired conspecific interactions. Neurochemical analysis revealed a marked increase in serotonin (5-HT) levels, whereas lipid peroxidation (MDA) remained unchanged, suggesting the absence of overt oxidative damage. Gene expression profiling demonstrated a dysregulated antioxidant response, suppression of apoptotic signaling, and pronounced upregulation of serotonergic receptors and transporters. To resolve system-level mechanisms, protein-protein interaction (PPI) network analysis identified BDNF and CREB1 as dominant regulatory hubs, with the serotonergic synapse pathway as the most significantly enriched term. Molecular docking further demonstrated direct binding of BPS to multiple serotonergic targets, including HTR1A and TPH2, supporting receptor-level interference. Expanded network and pathway analyses revealed coordinated enrichment of monoamine GPCR, oxidative stress, and inflammatory pathways. These findings demonstrate that BPS induces serotonergic dysregulation and network-level reprogramming rather than significant oxidative damage, leading to behavioural impairment. This study provides a multi-scale mechanistic framework linking molecular perturbations to neurobehavioural outcomes, identifying serotonergic signaling and BDNF-CREB1 pathways as central targets of BPS neurotoxicity.

## 1. Introduction

Bisphenol S (BPS; bis-(4-hydroxyphenyl)-sulfone) has emerged as one of the most widely used substitutes for bisphenol A (BPA) in the production of “BPA-free” consumer products, including thermal papers, food packaging materials, epoxy resins, and plastics (Manzoor et al., 2022; Yang et al., 2019). The increasing global restrictions on BPA due to its well-documented endocrine-disrupting effects have accelerated the adoption of BPS as an alternative. However, growing evidence indicates that BPS is environmentally persistent and biologically active, raising concerns regarding its safety. BPS has been detected in surface waters, sediments, and wastewater at concentrations ranging from trace levels to over 60 µg/L, indicating continuous exposure risks for aquatic organisms (Fabrello & Matozzo, 2022; Li et al., 2024; Yamazaki et al., 2015). Furthermore, widespread human exposure has been reported, with BPS detected in the majority of urine and serum samples across multiple populations (Liao et al., 2012; Wan et al., 2018).

Structurally similar to 17β-estradiol, BPS can bind to estrogen receptors and interfere with endocrine signaling pathways, exhibiting estrogenic, anti-androgenic, and thyroid-disrupting properties (Gupta & Haldar, 2017; Molina-Molina et al., 2013; Rosenmai et al., 2014). Consequently, BPS exposure has been associated with a range of adverse biological outcomes, including reproductive dysfunction, cytotoxicity, genotoxicity, and developmental abnormalities in various animal models (Ji et al., 2013; Moreman et al., 2017; Qiu et al., 2016). While the endocrine-disrupting effects of BPS are increasingly recognized, its impact on the central nervous system (CNS) and behaviour remains comparatively underexplored. Emerging evidence suggests that bisphenol analogues can interfere with neuroendocrine processes that regulate cognition, stress responses, and social behaviour (Saili et al., 2012; Weber et al., 2015), yet the underlying mechanisms remain poorly understood.

Among the proposed mechanisms of BPS-induced neurotoxicity, oxidative stress has been identified as a critical pathway. BPS exposure has been shown to increase the production of reactive oxygen species (ROS), disrupt antioxidant defense systems, and induce redox imbalance in multiple tissues, including neural tissue (Lee et al., 2008; Maćczak et al., 2017). In aquatic organisms, including zebrafish, BPS exposure has been associated with increased oxidative stress, lipid peroxidation, and altered antioxidant enzyme activity (Qiu et al., 2016; Salahinejad et al., 2021). Oxidative stress plays a fundamental role in neuronal function, as a delicate balance between ROS production and antioxidant capacity is essential for neurotransmission, synaptic plasticity, and neuronal survival (Ji, 2002; Kamat et al., 2016). Disruption of this balance can lead to functional alterations in neural circuits, even in the absence of overt cellular damage. Moreover, oxidative stress is closely linked to apoptosis, and perturbations in apoptotic signaling pathways such as those involving *p53, bax, bcl-2,* and *caspase-3* can influence neuronal survival and neurodevelopment (Elmore, 2007). Thus, the interaction between oxidative stress and apoptosis represents a key axis in understanding chemical-induced neurotoxicity.

In addition to redox-mediated mechanisms, neurotransmitter systems play a pivotal role in mediating behavioural outcomes following toxicant exposure. The serotonergic system is particularly important in this regard, as serotonin (5-hydroxytryptamine, 5-HT) regulates a wide range of behaviours, including mood, anxiety, social interaction, and cognition (Bacqué-Cazenave et al., 2020). During early development, serotonin also functions as a neurotrophic factor, influencing neuronal differentiation, migration, and synaptogenesis (Whitaker-Azmitia, 2001). Disruption of serotonergic signaling during critical developmental windows can therefore have long-lasting consequences on brain function and behaviour. Brain-derived neurotrophic factor (BDNF) and cAMP response element-binding protein 1 (CREB1) are key regulators of neuronal development, synaptic plasticity, learning, and social behaviour (Benito & Barco, 2015; Lu et al., 2013). BDNF and CREB1 function within interconnected signaling pathways that support neuronal survival and activity-dependent gene expression, and alterations in these pathways have been linked to neurodevelopmental and behavioural disorders (Carlezon et al., 2005; Cunha et al., 2010). Previous studies have demonstrated that environmental contaminants, including bisphenol analogues, can alter BDNF-CREB signaling, serotonin synthesis, receptor expression, and transporter activity, leading to dysregulated neurotransmission and behavioural abnormalities (Hasan et al., 2026; Narasimhamurthy et al., 2022; Semeão et al., 2025).

Zebrafish *(Danio rerio)* represent a well-established vertebrate model for investigating neurobehavioral toxicity due to their conserved neurobiological pathways, rapid development, and robust behavioural repertoire (Firdous et al., 2024). Social behaviour in zebrafish, including group preference and conspecific interactions, is particularly sensitive to disruptions in serotonergic signaling and serves as a reliable indicator of neurodevelopmental perturbation (Attaran et al., 2021; Salahinejad et al., 2020). These behaviours are critical for survival, as they influence predator avoidance, foraging efficiency, and reproductive success (Miller & Gerlai, 2012). Importantly, early-life exposure to environmental contaminants can lead to persistent alterations in these behaviours, even in the absence of overt morphological defects.

Despite increasing evidence of BPS toxicity, significant gaps remain in understanding how developmental exposure to environmentally relevant concentrations disrupts social behaviour and associated underlying molecular mechanisms. In particular, the integrated roles of oxidative stress, apoptotic signaling, serotonergic dysregulation, and network-level molecular interactions in shaping behavioural outcomes have not been comprehensively resolved. Addressing this gap is essential for advancing mechanistic insight into BPS-induced neurodevelopmental toxicity and its ecological relevance. Therefore, the present study investigated whether developmental exposure to an environmentally relevant concentration of BPS (30 µg/L) from 4 to 120 hours post-fertilization induces alterations in social behaviour in zebrafish larvae. To elucidate the underlying mechanisms, we combined behavioural assays with biochemical (serotonin and lipid peroxidation), molecular (RT-qPCR), and systems-level computational analyses, including protein-protein interaction (PPI) network modelling, pathway enrichment, and molecular docking. Specifically, we tested the hypothesis that BPS disrupts neurodevelopment through serotonergic pathway dysregulation coupled with oxidative stress-mediated signaling and network reorganization involving the neuroplasticity regulators BDNF and CREB1. By integrating experimental and computational approaches, this study provides a multi-scale mechanistic framework linking molecular perturbations to behavioural deficits, offering new insight into how BPS interferes with neurodevelopment and social function in aquatic organisms.

## 2. Materials and Methods

### 2.1. Animals

Zebrafish of the standard wild-type strain (AB strain) were sourced from the University of Saskatchewan’s Collaborative Science and Research Building (CSRB) vivarium facility. The fish were maintained in a recirculating aquatic system under a photoperiod of 14 hours light and 10 hours darkness. Environmental conditions were carefully controlled to maintain water temperature between 26 and 28°C, a pH range of 7.4 to 7.6, conductivity between 650 and 750 μS/cm, a total hardness of approximately 150 mg/L, and an alkalinity level of 120 mg/L expressed as calcium carbonate equivalents. Feeding occurred twice per day throughout the maintenance period. For breeding purposes, tanks were prepared by combining two female and one male fish, which were left overnight to allow natural spawning. Embryos were collected the following morning. The study received ethical approval from the University of Saskatchewan’s Animal Research Ethics Committee under permit number AUP20230068.

### 2.2. Chemicals

BPS with a minimum purity level of 99% was procured from Sigma Aldrich (Canada). The chemical compound was dissolved in dimethyl sulfoxide (DMSO) to generate stock solutions, which were subsequently stored at 4°C for use in subsequent experiments. During all experimental procedures, the concentration of DMSO solvent was controlled to remain at 0.01% (v/v) or lower. The accuracy of the measured concentration of BPS in the exposure medium was verified and reported elsewhere (Hasan et al., 2026). Please see supplementary file.

### 2.3 Experimental Design

Embryos at the four-hour post-fertilization (4-hpf) stage with chorion intact were randomly distributed into treatment groups and subjected to two experimental conditions: a vehicle control containing DMSO and a BPS-exposed group at a concentration of 30 μg/L. Both treatment groups were maintained in 100-mm petri dishes filled with 50 ml of E3 medium (5 mM NaCl, 0.7 mM KCl, 0.33 mM CaCl₂, and 0.33 mM MgSO₄) from 4-hpf through 120 hpf. Following the initial exposure period, larvae were transferred to 3.5 L rearing tanks at 6-dpf and reared through the completion of the study at 24-dpf. Following yolk sac absorption, zebrafish larvae were initially fed GEMMA Micro 75 as a starter diet twice daily for the first three days. Thereafter, larvae were fed Artemia twice daily for the remainder of the experimental period. At 21-dpf, social behavior was assessed using standardized behavioral tests to evaluate social preference and interaction patterns. At 24-dpf, larvae were euthanized and head tissues were collected for subsequent molecular analysis (including gene expression profiling) and biochemical assays. A total of six independent breeding events were used in this study. For each breeding event, 50 embryos were collected at 4 hpf and randomly allocated to either the control or BPS treatment group (25 embryos per treatment). Thus, each treatment consisted of six biological replicates derived from the same six breeding events.

### 2.4. Social preference test

At 21 dpf, social behaviour was evaluated using a custom-built three-chambered apparatus measuring 24 cm (length) × 6 cm (width) × 6 cm (height), constructed from 2 mm sheet material. The apparatus consisted of a central compartment flanked by two lateral zones: one designated as the empty zone and the other as the social zone. Five conspecific stimulus larvae, sourced from the same housing tank, were placed in the social zone prior to testing. The focal larva was then introduced into the central chamber and allowed freely navigate the entire apparatus over a 10-minute trial period. The percentage of time the focal fish spent in the conspecific zone, defined as the area adjacent to the conspecific box, was calculated as an indicator of social preference. To minimize positional bias, the location of the conspecific box (left or right side) was alternated between replicates (n= 28) for each treatment group.

### 2.5 Serotonin level assessment

Total serotonin (5-hydroxytryptamine, 5-HT) levels were quantified in the head of zebrafish homogenates using a serotonin-specific ELISA kit (Biohippo) following the manufacturer’s protocol. A total of 2-3 heads were pooled together per biological replicate (n = 6) and homogenized in ice-cold phosphate-buffered saline (PBS; approximately 50 mg tissue in 450 µL PBS). Homogenates were centrifuged at 5000 × g for 5 min, and the supernatants were collected for analysis. Briefly, 50 µL of each sample was added to the ELISA plate, followed by biotin-detection antibody working solution, and incubated at 37 °C for 45 min. After washing, HRP-streptavidin conjugate working solution was added and incubated at 37 °C for 30 min. The wells were washed again, incubated with TMB substrate for 20 min at 37 °C, and the reaction was stopped with stop solution. Absorbance was measured at 450 nm.

### 2.6. Malondialdehyde (MDA) content

Lipid peroxidation in head tissues of 28 dpf zebrafish larvae, expressed as malondialdehyde (MDA) content, was quantified using the Lipid Hydroperoxide Assay Kit (ab133085, Abcam) according to the manufacturer’s instructions. For each treatment group, six biological replicates were analyzed, with each replicate consisting of a pooled sample of 2-3 larval heads. Samples were homogenized on ice in HPLC-grade water, followed by extraction of lipid hydroperoxides into chloroform using the kit-provided Extract R reagent. This step minimizes interference from hydrogen peroxide and endogenous ferric ions. For the assay, 500 μL of the chloroform extract was combined with 450 μL chloroform:methanol (2:1, v/v) and 50 μL of freshly prepared chromogen solution prepared by mixing equal volumes of FTS Reagent 1 and FTS Reagent 2. After incubation for 5 min at room temperature, absorbance was measured at 500 nm using a 96-well glass plate. Lipid hydroperoxide concentrations were calculated from a 13-HpODE standard curve and expressed as μM.

### 2.7. Gene expression analysis

Gene expression in 28 dpf zebrafish larvae head tissue was assessed by real-time quantitative polymerase chain reaction (RT-qPCR) targeting three functional gene categories: antioxidant markers (*cat, gpx, mn/sod, cu-zn/sod*), apoptotic pathway genes (*p53, bcl2, bax, casp-3*), and serotonergic system genes (*tph2, htr1aa, htr1ab, htr1b, htr1d, htr2aa, htr2b, slc6a4a, slc6a4b*). Total RNA was extracted from pooled head tissue (six replicates; 2-3 head per replicate) using a commercial purification kit (RNeasy Mini Kit, Qiagen) including DNase treatment, and RNA quality was verified spectrophotometrically (A260/A280 ratio). Complementary DNA was synthesized from 1 μg total RNA using reverse transcription, and RT-qPCR was performed in triplicate using SYBR Green master mix with 0.8 μL each of forward and reverse primers and 2 μL cDNA template in 20 μL reactions. Gene expression was normalized to the reference gene *β-actin*, PCR efficiency was optimized to exceed 95%, and relative expression levels were calculated using the 2-ΔΔCt method. Primer sequences are provided in Table S7 (Supplementary Materials).

### 2.8 Protein-protein interaction network analysis

A PPI network was constructed from the human orthologs of BPS-regulated zebrafish genes identified by RT-qPCR (n=19 seed nodes) supplemented with established interaction partners from the literature (n=26 curated edges). Protein interaction data were retrieved from the STRING database (v12.0; https://string-db.org; Szklarczyk et al., 2023) using a combined interaction score threshold of ≥ 0.400, returning 71 high-confidence interactions. The network was constructed and analyzed using NetworkX v3.2 (Python 3.11). Four centrality metrics were calculated for each node: degree centrality, betweenness centrality, closeness centrality, and eigenvector centrality. A composite hub score was computed as the arithmetic mean of the four normalized centrality values. Pathway enrichment analysis was performed via the STRING enrichment API against KEGG, Reactome, WikiPathways, and GO Biological Process databases. The significance threshold for enrichment was FDR < 0.05 (Benjamini-Hochberg correction). The network was visualized using Matplotlib v3.8 and exported in Cytoscape-compatible JSON format. Hub centrality scores and network topology metrics for the 19 seed genes are summarized in Table 1 (Supplementary Data). The PPI network and centrality heatmap are illustrated in Figures 6 A,B,C.

### 2.9 Molecular docking

The three-dimensional structure of BPS (bisphenol S; PubChem CID: 6623; molecular formula: C12H10O4S; MW: 250.27 g/mol; SMILES: O=S(=O)(c1ccc(O)cc1)c1ccc(O)cc1) was retrieved from PubChem in SDF format with optimized 3D coordinates. Ligand PDBQT preparation was performed using OpenBabel v3.1.0 with Gasteiger partial charge assignment and all-hydrogen addition. Six neurological and endocrine receptor targets were selected based on their relevance to BPS-regulated pathways: 5-HT1A receptor (PDB: 7E2Y), 5-HT2B receptor (PDB: 4IB4), 5-HT2A receptor (PDB: 6WGT), serotonin transporter SERT/SLC6A4 (PDB: 6DZV), tryptophan hydroxylase 2 TPH2 (PDB: 4V04), and estrogen receptor alpha ERα (PDB: 1A52). Receptor PDB structures were downloaded from the RCSB Protein Data Bank and prepared as rigid PDBQT files using OpenBabel with polar-hydrogen addition. Non-standard atom types arising from crystallographic conditions (e.g., Au from crystal contacts in 1A52) were removed prior to docking. Binding site centroids were computed from co-crystallized ligand HETATM atom coordinates (centroid method) or catalytic residue anchor atoms where co-crystal ligands were absent. Docking was performed with AutoDock Vina v1.2.5 (Eberhardt et al., 2021) with grid boxes of 25 × 25 × 25 Å centered on each active site, exhaustiveness = 8, and 9 binding modes generated per run. The best-ranked pose (mode 1) affinity was used for comparison. Binding affinity ≤ -5.0 kcal/mol was considered indicative of meaningful interaction, consistent with established thresholds in zebrafish endocrine disruptor docking studies (Shan et al., 2019). Predicted binding affinities (kcal/mol) for BPS against all six targets are summarized in Table S2 (Supplementary Data).

### 2.10 Computational novel target analysis

The Comparative Toxicogenomics Database (CTD; https://ctdbase.org; Davis et al., 2021) was queried for all gene-BPS interaction records in fish and mammalian systems. Genes with a CTD interaction count ≥ 3 were retained as high-confidence targets (n = 61), ensuring inclusion of only the most reproducible gene-chemical associations. Resolved gene identifiers were submitted to the STRING database (Danio rerio, taxon 7955; score ≥ 0.4), returning 312 interactions. K-means clustering (k = 3) was applied to the four-dimensional centrality feature space (scikit-learn v1.4) to partition the network into functionally coherent modules. Functional protein class annotation was performed manually guided by UniProt and CTD functional categories, mirroring the classification scheme of dos Santos et al. (2022) for BPF. Pathway enrichment was performed via the STRING enrichment API (KEGG, Reactome, WikiPathways, GO Biological Process; FDR < 0.05). Network figures were generated using Matplotlib v3.8 and NetworkX v3.2. Hub centrality scores for the 61-gene zebrafish network, pathway enrichment results, and functional class distributions are summarised in Tables 4-6 (Supplementary Data). Network topology, hub gene rankings, and enriched pathways are illustrated in Figures 7-8.

### 2.11. Statistical analysis

All statistical analyses were performed in Python (version 3.14.0) using the following libraries: SciPy (v1.17.1), NumPy (v2.4.4), pandas (v3.0.2), matplotlib (v3.10.8), and seaborn (v0.13.2). Data are presented as mean ± standard error (SE) with 95% confidence intervals. Prior to group comparisons, normality was assessed using the Shapiro-Wilk test (scipy.stats.shapiro). As most datasets deviated from a normal distribution (p < 0.05), differences between the BPS-exposed and control groups were evaluated using the two-tailed Mann-Whitney U test (scipy.stats.mannwhitneyu). The significance threshold was set at α = 0.05 for all analyses. All figures were generated using matplotlib and seaborn.

## 3. Results

### 3.1 Effect of BPS on Social Behaviour

Developmental exposure to BPS significantly affected social behaviour in zebrafish larvae. Time spent in the conspecific zone was significantly reduced in the BPS-exposed group compared with the control group (Mann-Whitney U test, U = 287, p = 0.0154; two-tailed; Fig. 1). The BPS exposed larvae spent approximately 52.56% of the time in the conspecific zone (median, n = 30) compared to 74.26% in the control group (median, n = 30), representing a 21.71% reduction in social proximity and engagement. This difference was accompanied by a medium effect size (Cohen’s d = 0.6428), indicating a biologically meaningful impairment in social behaviour. The Hodges-Lehmann point estimate of the difference between medians was -14.77, further demonstrating a substantial shift in social behaviour following BPS exposure. These findings suggest that developmental BPS exposure impairs normal social interactions and proximity-seeking behaviour in zebrafish larvae, with detectable consequences for conspecific engagement.

**Figure 1.**
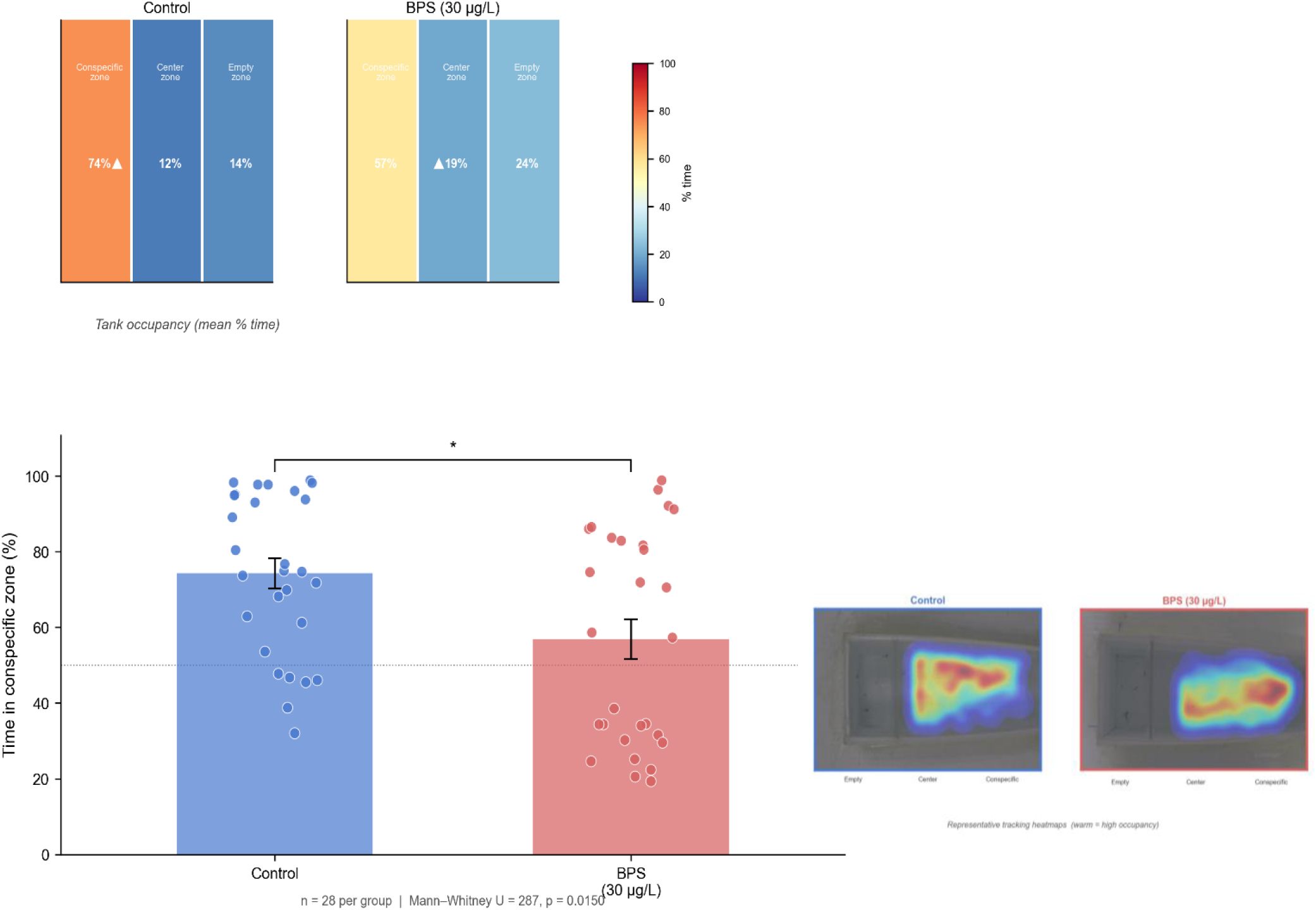
Developmental BPS exposure impairs social preference behaviour in zebrafish larvae at 21 dpf. (A) Tank occupancy heatmaps showing the mean percentage of time spent in each zone (conspecific, center, and empty) for control and BPS-exposed (30 µg/L) larvae. Control fish exhibited a strong preference for the conspecific zone (74%), whereas BPS-exposed larvae showed a reduced preference (57%) and increased time in the center (19%) and empty zones (24%), indicating altered social exploration patterns. (B) Quantification of social preference, expressed as percentage of time spent in the conspecific zone. Each point represents an individual larva (n = 28 per group), with bars indicating mean ± SE. Developmental exposure to BPS significantly reduced time spent near conspecifics compared to controls, demonstrating impaired social affiliation. The dashed horizontal line in (B) indicates equal distribution across zones (50%), representing the absence of social preference. (C) Representative tracking heatmaps illustrating spatial occupancy of individual larvae during the assay. Warmer colors indicate higher occupancy. Control larvae showed concentrated activity near the conspecific zone, whereas BPS-exposed larvae displayed more dispersed movement with reduced localization near conspecifics.

**Figure 2.**
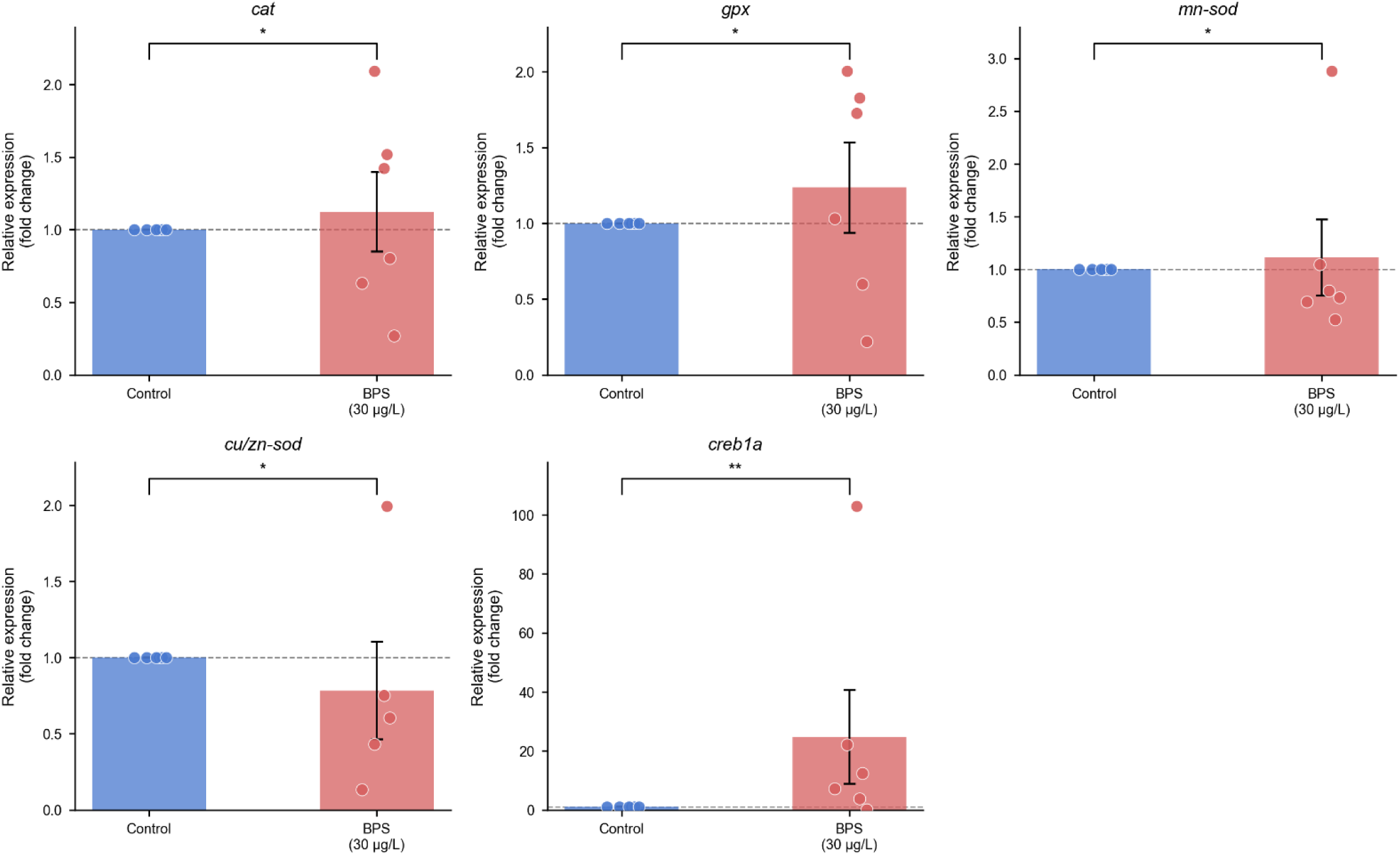
Developmental BPS exposure disrupts antioxidant gene expression in zebrafish larvae at 28 dpf. Data are presented as mean ± SE (n = 6 biological replicates; each replicate consists of pooled head tissue from 2-3 larvae). Individual data points are shown. Statistical significance was determined using the Mann-Whitney U test (*p < 0.05, **p < 0.01). The dashed horizontal line represents control-normalized expression (fold change = 1).

**Figure 3.**
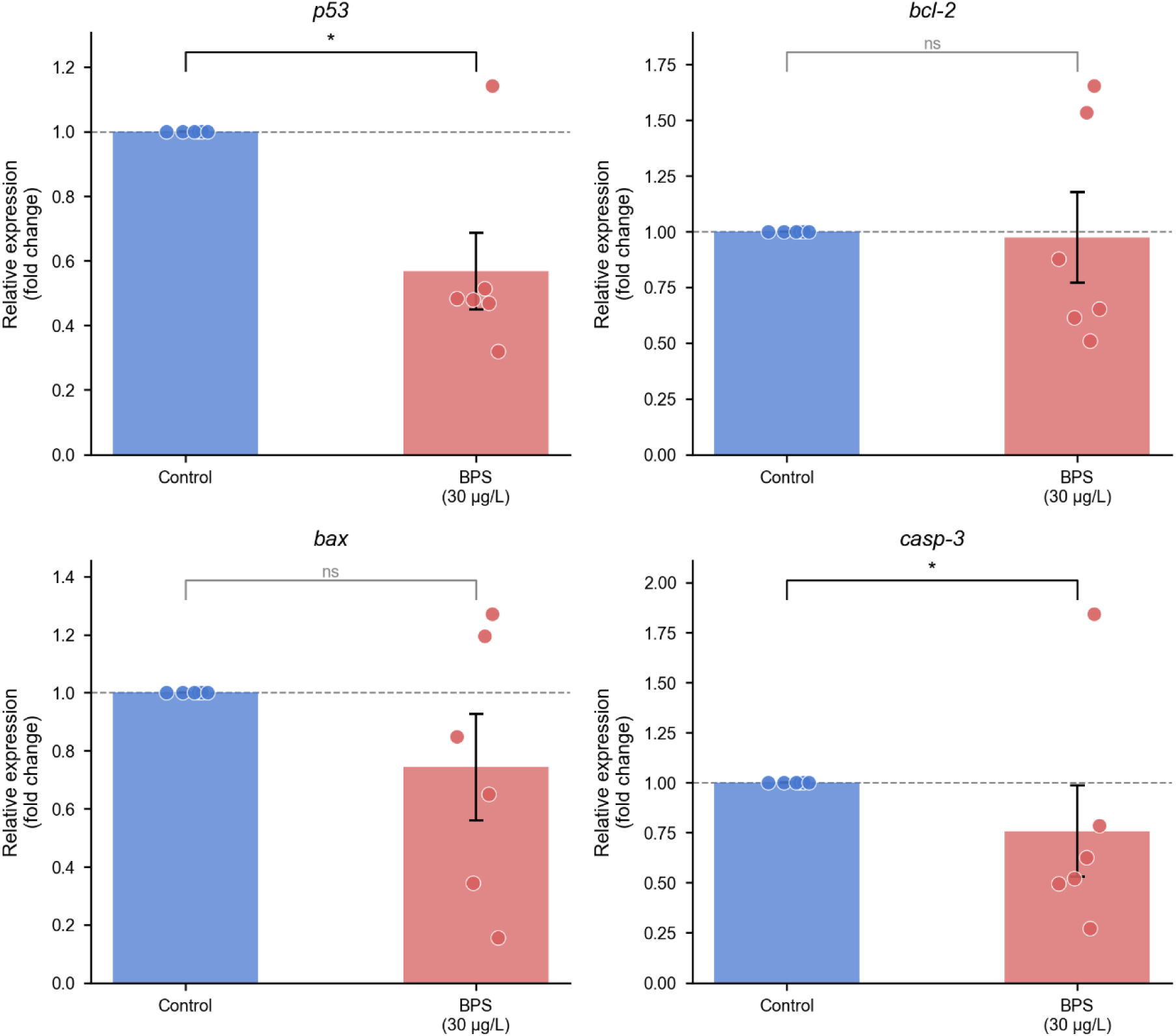
Developmental BPS exposure suppresses apoptotic signaling pathways in zebrafish larvae. Data are presented as mean ± SE (n = 6 biological replicates; pooled head tissue), with individual data points overlaid. Gene expression levels were normalized to β-actin using the 2⁻ΔΔCt method, and the dashed horizontal line represents control-normalized expression (fold change = 1). Statistical significance was determined using the Mann–Whitney U test (*p < 0.05; ns = not significant).

**Figure 4.**
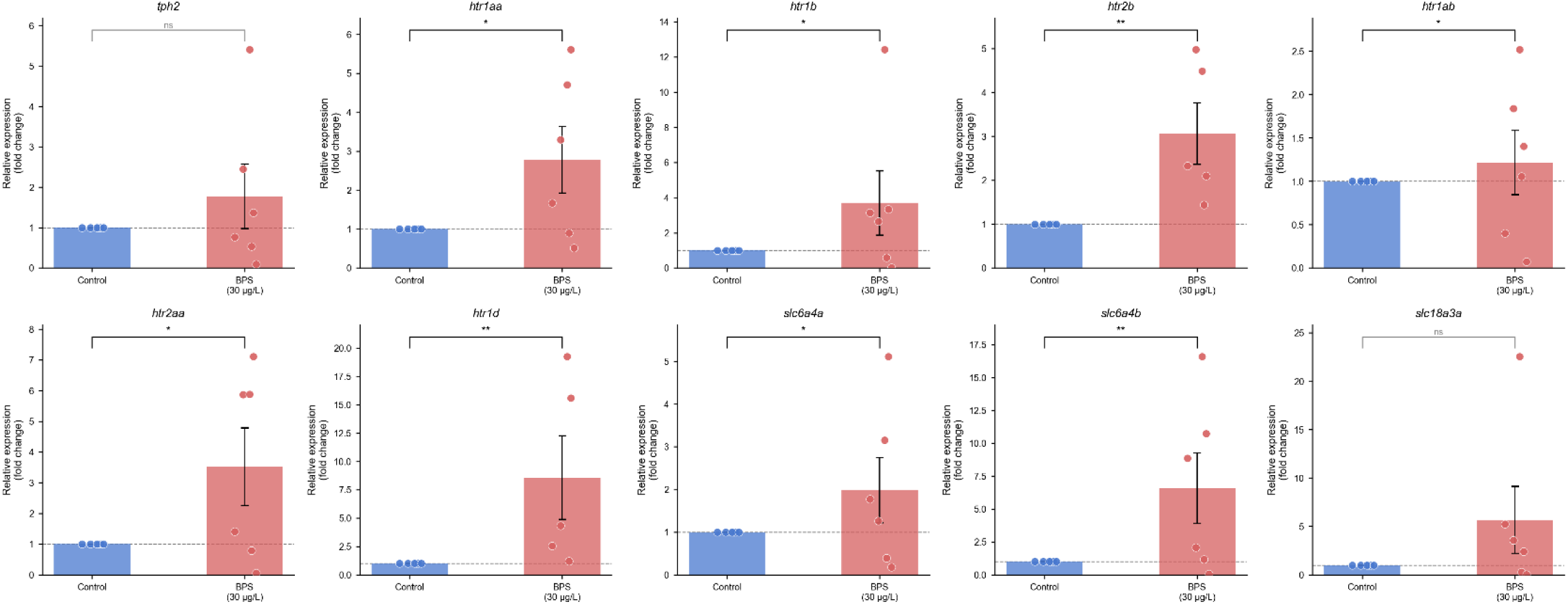
Developmental BPS exposure induces widespread upregulation of serotonergic signaling pathways in zebrafish larvae. Data are presented as mean ± SE (n = 6 biological replicates; pooled head tissue), with individual data points shown. Gene expression levels were normalized to β-actin using the 2⁻ΔΔCt method, and the dashed horizontal line indicates control-normalized expression (fold change = 1). Statistical significance was determined using the Mann-Whitney U test (*p < 0.05, **p < 0.01; ns = not significant).

### 3.2 Effects of BPS exposure on lipid peroxidation

Developmental exposure to BPS did not significantly affect malondialdehyde (MDA) levels in 28 dpf zebrafish larvae (Mann-Whitney U test, p = 0.3095; Fig. 5B). These findings indicate that BPS exposure did not induce detectable lipid peroxidation under the experimental conditions tested.

**Figure 5.**
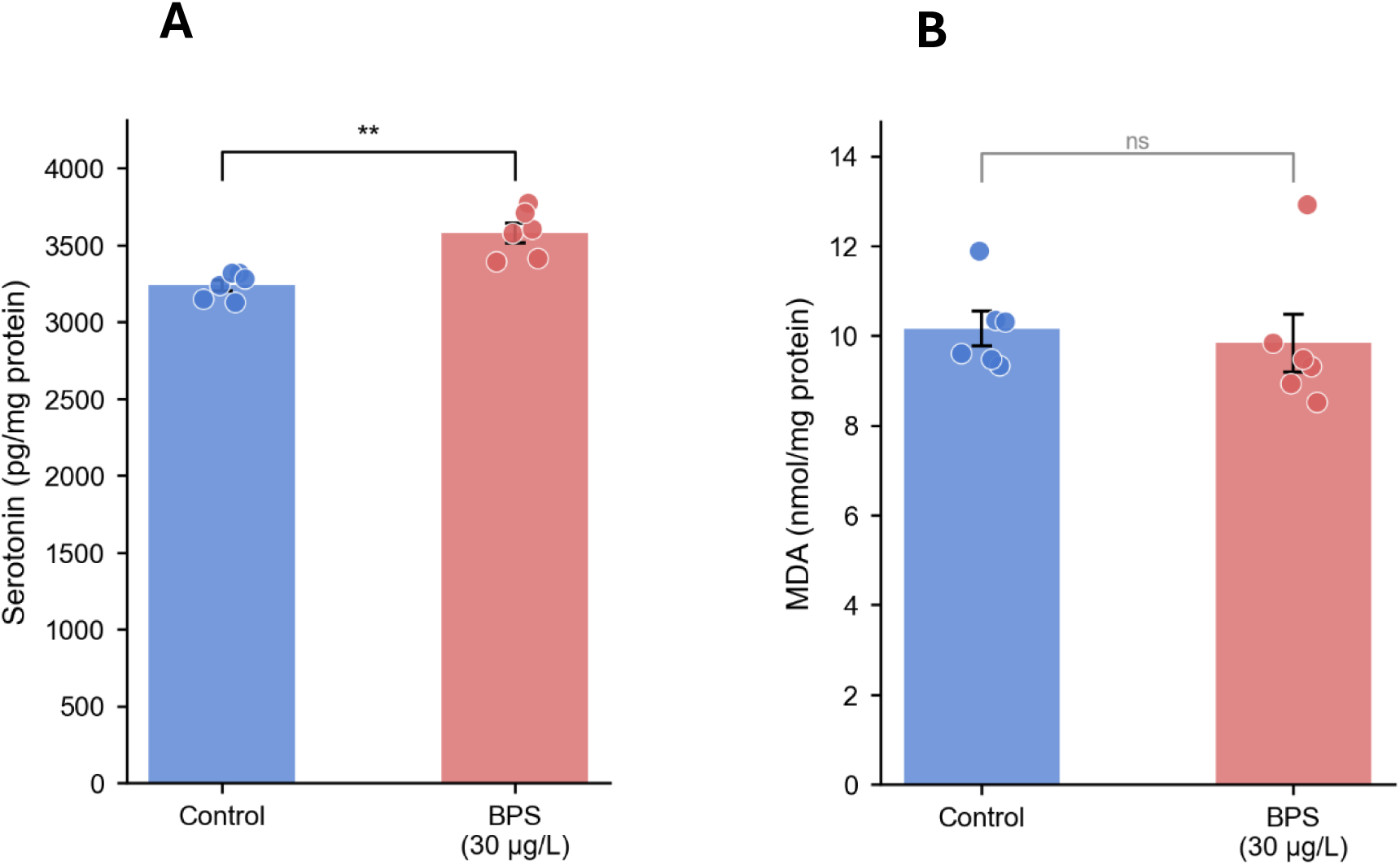
Developmental BPS exposure elevates serotonin levels without inducing lipid peroxidation in zebrafish larvae. Data are presented as mean ± SE (n = 6 biological replicates; pooled head tissue), with individual data points shown. Serotonin (A) and MDA (B) levels were quantified using ELISA and lipid hydroperoxide assays, respectively. Statistical significance was determined using the Mann-Whitney U test (**p < 0.01; ns = not significant).

### 3.3 Effect of BPS on Serotonin (5-HT) Levels

Developmental exposure to BPS significantly elevated serotonin (5-HT) levels in zebrafish larvae. The control group exhibited a mean 5-HT level of 3239.98 ± 84.33 pg/mg protein (median: 3260.39, n = 6; 95% CI: 3172.50-3307.45), whereas the BPS-exposed group showed a substantially higher mean of 3579.86 ± 154.74 pg/mg protein (median: 3592.64, n = 6; 95% CI: 3456.04-3703.67). This represents a considerable elevation of

339.88 pg/mg protein (10.49% increase) in mean 5-HT levels, with a median difference of 332.25 pg/mg protein. The Mann-Whitney U test revealed a highly statistically significant difference between treatment groups (U = 0, p = 0.0022, two-tailed, exact p-value). The effect size was large (Cohen’s d = 2.7276), indicating a substantial and biologically meaningful difference between groups. The Hodges-Lehmann point estimate of the difference between medians was 332.25, providing a robust, distribution-free estimate of the median difference. These findings demonstrate that developmental BPS exposure significantly increases serotonergic signalling in zebrafish larvae.

### 3.4 Gene expression changes

Exposure to 30 µg/L BPS significantly altered the expression of genes associated with oxidative stress, apoptosis, and serotonergic signaling in zebrafish larvae. With respect to antioxidant defense, catalase (*cat*) and *gpx* were significantly upregulated by 1.7-fold (p = 0.0316) and 1.6-fold (p = 0.0283), respectively. In contrast, both *mn-sod* and *cu/zn-sod* were significantly downregulated by 0.7-fold (p = 0.0476) and 0.8-fold (p = 0.0476), respectively, while *creb1a* expression was markedly upregulated (∼30-fold; p = 0.0079), indicating an unbalanced antioxidant response.

In the apoptotic pathway, *caspase-3* and *p53* were significantly downregulated by 0.7-fold (p = 0.0152) and 0.6-fold (p = 0.0152), respectively. Meanwhile, *bax* showed a non-significant decreasing trend (0.7-fold; p = 0.0606), and *bcl-2* expression remained unchanged (p = 0.0606), suggesting suppression of key pro-apoptotic signaling components.

In the serotonergic signaling pathway, the serotonin synthesis gene *tph2* exhibited an increasing trend (∼3.0-fold), although this change was not statistically significant (p = 0.1000). In contrast, multiple serotonin receptor genes were significantly upregulated, including *htr1aa* (∼3.2-fold; p = 0.0358), *htr1ab* (∼1.4-fold; p = 0.0476), *htr1b* (∼4.5-fold; p = 0.0476), *htr1d* (∼8.5-fold; p = 0.0079), *htr2aa* (∼4.2-fold; p = 0.0476), and *htr2b* (∼3.0-fold; p = 0.0079). Additionally, serotonin transporter genes were significantly upregulated, with *slc6a4a* increasing by ∼2.8-fold (p = 0.0286) and *slc6a4b* by ∼8.0-fold (p = 0.0079). These findings demonstrate that BPS exposure disrupts redox homeostasis, suppresses apoptotic signaling, and induces a pronounced upregulation of serotonergic receptor and transporter systems, indicating coordinated alterations in neurodevelopmental and neurotransmission-related pathways.

### 3.5 PPI Network Analysis Identifies BDNF and CREB1 as Central Hubs

The full hub-score rankings, together with degree, betweenness, closeness, and eigenvector centrality values for all 19 genes, are provided in Table S1. The network graph and centrality heatmap are shown in Figures 6-8. K-means clustering (k = 3) partitioned the network into a serotonin receptor cluster (HTR1A, HTR2A, HTR2B, HTR3A, HTR4, HTR7), a neurotrophin/stress hub cluster dominated by BDNF and CREB1, and a monoamine metabolism cluster (MAOA, MAOB, TPH2). Pathway enrichment of the full STRING network (Table S3) identified the serotonergic synapse (KEGG; FDR = 1.04 × 10⁻¹³) as the most significantly enriched canonical pathway, consistent with the experimental gene-expression findings reported above.

**Figure 6.**
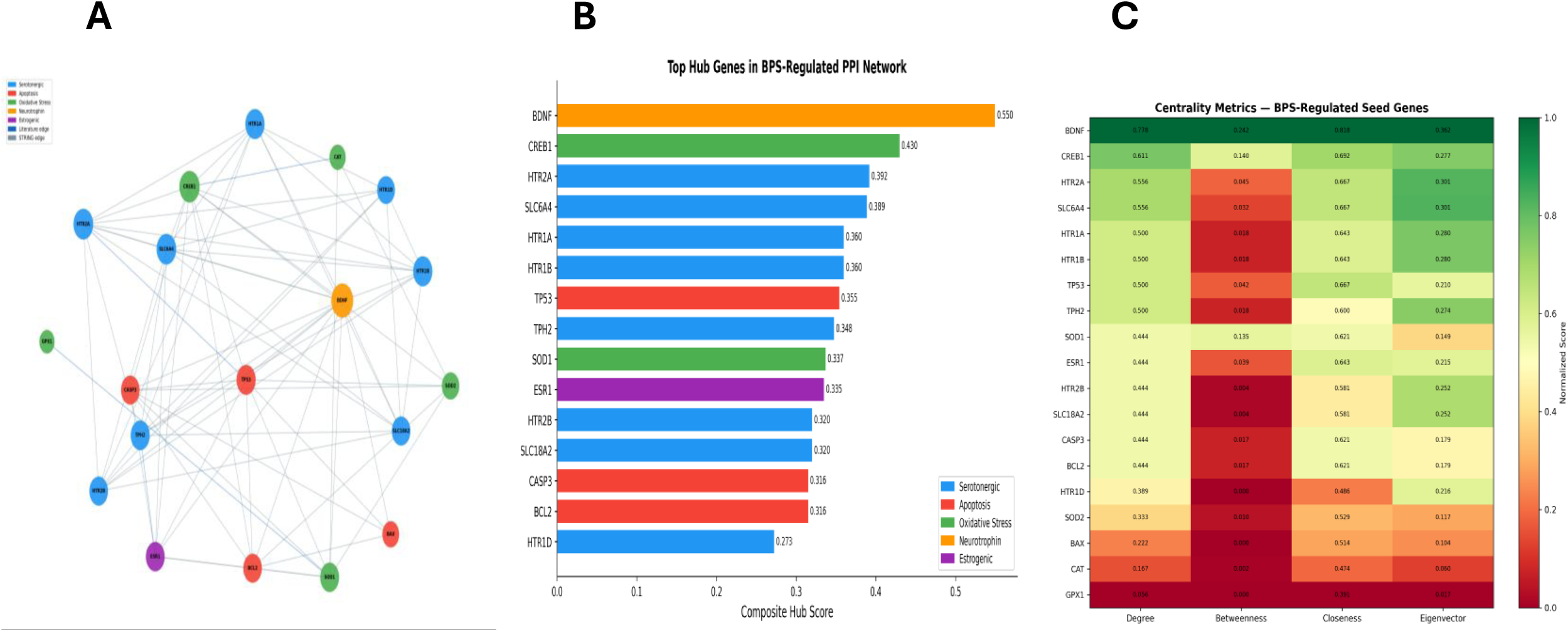
Integrated protein-protein interaction network analysis identifies key hub regulators of BPS-induced neurotoxicity. (a) The PPI network constructed from BPS-regulated genes (human orthologs; STRING v12.0) reveals a highly interconnected interactome (19 seed nodes, 75 edges), with node size proportional to composite hub score and colors indicating functional categories (serotonergic, oxidative stress, apoptosis, neurotrophic, and estrogenic pathways). (b) Ranking of top hub genes based on composite centrality (mean of degree, betweenness, closeness, and eigenvector centrality) identifies BDNF and CREB1 as dominant hubs, followed by key serotonergic components (*HTR2A, SLC6A4, HTR1A/B*) and apoptosis-related regulators (*TP53, CASP3*), highlighting cross-pathway integration. (c) Centrality heatmap (z-score normalized) demonstrates that BDNF and CREB1 exhibit consistently high scores across all centrality metrics, whereas serotonergic receptor genes show high degree but comparatively lower betweenness, indicating strong local connectivity but limited global network control. Collectively, this analysis reveals a BDNF-CREB1-centered regulatory axis linking serotonergic signaling, oxidative stress, and apoptosis, providing a systems-level framework for BPS-induced neurotoxicity.

**Figure 7.**
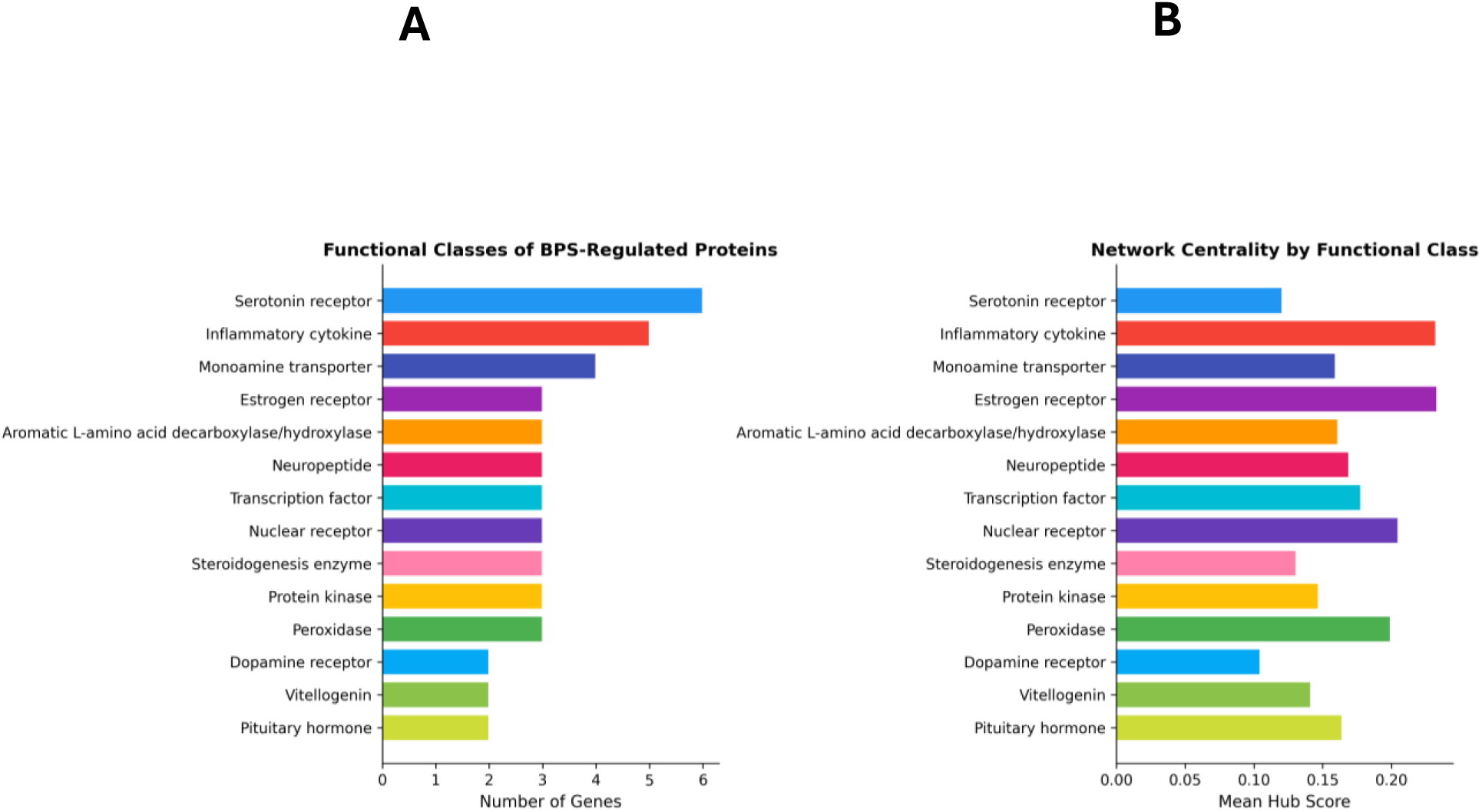
(A) Functional classification of BPS-regulated proteins shows enrichment of serotonin receptors, inflammatory cytokines, monoamine transporters, and estrogen receptors, alongside enzymes involved in neurotransmitter synthesis and oxidative stress regulation. (B) Network centrality analysis by functional class demonstrates that inflammatory cytokines and estrogen receptor pathways exhibit the highest mean hub scores, followed by monoaminergic and transcriptional regulators, whereas classical neurotransmitter receptor classes show moderate centrality.

**Figure 8.**
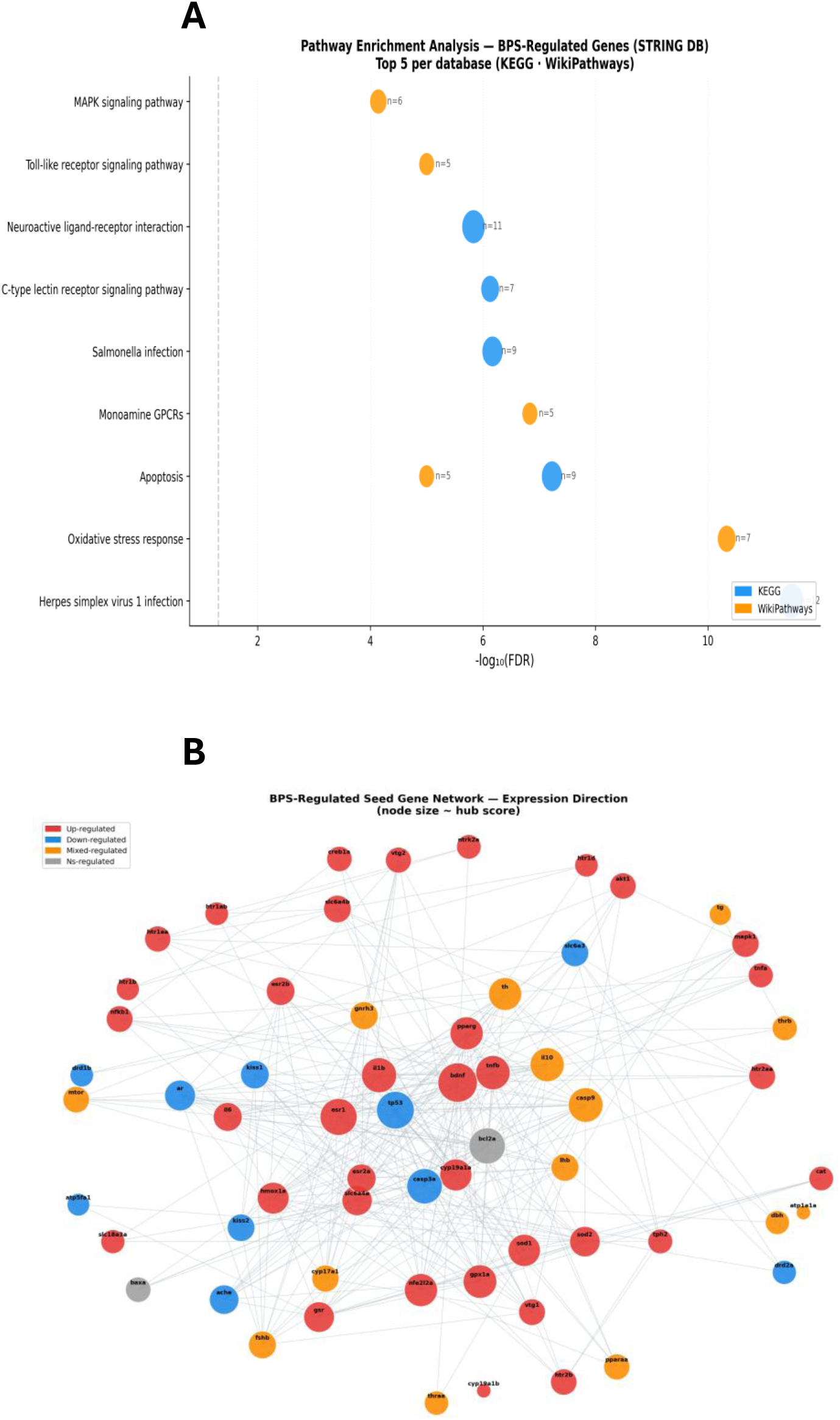
Integrated pathway enrichment and expression-resolved network analysis reveal coordinated modulation of neuroinflammatory and monoaminergic signaling by BPS. (a) Pathway enrichment analysis of BPS-regulated genes (STRING DB; top 5 pathways per database) highlights significant overrepresentation of neuroactive ligand-receptor interaction, monoamine GPCR signaling, MAPK signaling, Toll-like receptor signaling, apoptosis, and oxidative stress response pathways. KEGG pathways (blue) and WikiPathways (orange) show strong enrichment based on - log10(FDR), with bubble size proportional to gene count (n), indicating that both neurotransmission and immune-related signaling cascades are major targets of BPS exposure. (b) Expression-resolved protein-protein interaction network of BPS-regulated seed genes (node size proportional to hub score) illustrates the directionality of transcriptional responses, with upregulated genes (red), downregulated genes (blue), mixed-regulation nodes (orange), and non-significant genes (gray). The network displays a densely connected core centered on key regulators including *bdnf, tp53, il1b, tnfα*, and serotonergic components, with widespread upregulation across inflammatory and stress-response pathways, contrasted by selective downregulation of monoaminergic transporters and receptors.

### 3.6 Molecular Docking Demonstrates Direct BPS-Receptor Binding

Molecular docking of BPS against six neurological and endocrine receptor targets revealed binding affinities ranging from -2.706 to -6.274 kcal/mol (Table S2; Fig. 6). The strongest interaction was with the 5-HT1A receptor (HTR1A; -6.274 kcal/mol), followed by TPH2 (-5.805 kcal/mol), ERα (-5.778 kcal/mol), HTR2B (-5.675 kcal/mol), and HTR2A (-5.067 kcal/mol) all exceeding the -5.0 kcal/mol significance threshold. The serotonin transporter SERT showed weaker binding (-2.706 kcal/mol). These results provide direct computational evidence that BPS can occupy the orthosteric binding pockets of multiple serotonin receptors simultaneously, offering a mechanistic basis for the coordinated serotonergic gene expression changes observed experimentally. BPS binding at TPH2 further suggests potential interference with the rate-limiting step of serotonin biosynthesis, while ERα binding corroborates the established estrogenic activity of BPS and links it to the estrogen-serotonin regulatory axis.

### 3.7 Novel molecular targets of BPS-computational network analysis

Computational analysis of 61 high-confidence CTD-derived BPS-responsive genes (interaction count ≥ 3) in the zebrafish STRING network (taxon 7955) yielded a network of 61 nodes and 312 edges, with an average node degree of 10.23 and a PPI enrichment p-value < 1.0×10^-^¹⁶ (Table S4; Fig. 7). K-means clustering (k = 3) partitioned the network into three functional modules: Cluster 1 (n = 40 nodes) was dominated by serotonin receptors and monoamine transporters; Cluster 2 (n = 19 nodes) was enriched for inflammatory cytokines and apoptosis regulators; and Cluster 3 (n = 2) encompassed steroidogenesis enzymes.

Hub gene analysis of the expanded network again identified BDNF as the principal hub (hub score: 0.392; degree: 27), followed by TP53 (0.354), ESR1 (0.338), BCL2A (0.323), and CASP3A (0.305). Functional class analysis revealed that the BPS network encompasses 25 distinct protein functional classes, with serotonin receptors (n = 6), inflammatory cytokines (n = 5), monoamine transporters (n = 4), and aromatic L-amino acid decarboxylase/hydroxylase enzymes (n = 3) as the most represented classes. Pathway enrichment of the 61-gene network (Table S5) identified Apoptosis (KEGG; FDR = 5.94×10⁻⁸), Oxidative Stress Response (WikiPathways; FDR = 4.65×10⁻¹¹), Monoamine GPCRs (WikiPathways; FDR = 1.47×10⁻⁷), and Neuroactive ligand-receptor interaction (KEGG; FDR = 1.47×10⁻⁶) as the most significantly enriched terms.

## 4. Discussion

The present study demonstrates that developmental exposure to BPS at an environmentally relevant concentration (30 µg/L) induces significant impairment in social behaviour in zebrafish larvae, accompanied by pronounced dysregulation of serotonergic signaling and redox-related pathways. Notably, these behavioural deficits occurred in the absence of increased lipid peroxidation, indicating that BPS-induced neurotoxicity is not driven by overt oxidative damage but rather by neurochemical and transcriptional reprogramming. A key finding of this study is the significant elevation of serotonin levels, together with widespread upregulation of serotonergic receptors and transporters, highlighting a central role of serotonergic system disruption in mediating behavioural outcomes. Systems-level analyses further revealed that these molecular alterations are organized within a highly interconnected regulatory network, with BDNF (hub score: 0.550) and CREB1 (0.430) identified as dominant hubs and the serotonergic synapse pathway emerging as the most significantly enriched term (FDR=1.04 × 10⁻¹³). Importantly, molecular docking demonstrated that BPS can directly bind multiple serotonergic and endocrine targets, including HTR1A and TPH2, providing mechanistic support for receptor-level interference. Collectively, these findings indicate that BPS acts as a multi-target neuroactive compound, disrupting serotonergic neurotransmission and associated regulatory networks, ultimately leading to impaired social behaviour.

The toxicity of BPS in fish is frequently associated with oxidative stress (Shehna Mahim et al., 2021); however, the present findings indicate a more nuanced mechanism linking redox imbalance with neurochemical disruption. In the current study, zebrafish larvae exposed to 30 µg/L BPS exhibited a significant elevation in ROS, indicating the induction of an oxidative environment at the cellular level. This increase was accompanied by a selective and non-uniform transcriptional modulation of antioxidant defense genes, reflecting a disrupted rather than coordinated antioxidant response. Specifically, the upregulation of catalase (*cat*) and glutathione peroxidase *(gpx)*, in combination with the downregulation of *Mn-sod* and *Cu/zn-sod*, demonstrates a clear dysregulation within the hierarchical antioxidant defense system (Jomova et al., 2024). Given that SODs represent the primary line of defense by catalyzing the dismutation of superoxide radicals into hydrogen peroxide, their reduced expression suggests impaired upstream detoxification capacity, potentially leading to the intracellular accumulation of superoxide anions (Anwar et al., 2024). In contrast, the elevated expression of downstream enzymes such as catalase and GPx indicates a compensatory attempt to mitigate increased hydrogen peroxide and secondary ROS species (Ighodaro & Akinloye, 2018). This variance between upstream and downstream antioxidant components highlights a breakdown in redox homeostasis and suggests inefficient ROS clearance rather than a fully adaptive response. The simultaneous upregulation of *creb1a* further supports the activation of stress-responsive signaling pathways (Dworkin et al., 2007). As a transcription factor involved in neuronal survival, plasticity, and oxidative stress responses, increased *creb1a* expression likely reflects an adaptive neuroprotective mechanism aimed at maintaining cellular integrity under redox challenge (Dworkin et al., 2007). Overexpression of *creb1a* has been shown to confer neuroprotection against oxidative stress in the brain, as demonstrated in transgenic rodent studies. This suggests that the increased expression observed in the present study may represent an adaptive protective response to elevated oxidative stress, potentially contributing to the absence of lipid peroxidation (Lee et al., 2009). Wang et al., 2023 reported that early-life BPS exposure in zebrafish led to a significant reduction in brain lipid content and metabolic activity, suggesting that BPS-induced neurotoxicity may involve alternative mechanisms beyond classical oxidative damage (Wang et al., 2022). Despite increased oxidative stress study does not report increased classical apoptotic pathways. Significant downregulation of *caspase-3* and *p53*, with no observable changes in *bax* or *bcl-2* suggests a suppression or temporal uncoupling of intrinsic apoptotic signaling at 28 dpf. The absence of apoptotic activation, despite elevated ROS levels, indicates that BPS-induced stress may not directly induce cell death pathways but instead modulate cellular function through sublethal mechanisms. These findings support the notion that BPS-induced neurotoxicity is mediated not by disruption of redox-sensitive developmental and signaling processes, which may have downstream consequences for neuronal function and behaviour. Consistent with these observations, computational network analysis revealed CREB1 as the second-ranked hub gene (hub score: 0.430) directly linked to TP53 and BCL2 in the BPS interactome, while oxidative stress pathways (WikiPathways; FDR= 4.65 × 10^-^¹¹) and apoptosis (KEGG; FDR = 5.94 × 10^-^⁸) ranked among the top enriched terms. The network co-clustering of *casp3a* and *tp53* with antioxidant nodes further supports the notion of a sublethal, non-apoptotic stress response in which CREB1 acts as a transcriptional integrator rather than a trigger of classical cell-death programs.

The elevation in brain serotonin levels following developmental BPS exposure is noteworthy, as serotonin plays a critical role in regulating mood, anxiety, social behaviour, and cognitive function in vertebrates (Bacqué-Cazenave et al., 2020; Jayanthi & Ramamoorthy, 2025). The early ontogeny of the serotonergic system is important for neuronal growth and brain development, with 5-HT serving as a major neuromodulator that influences various behaviours through multiple linked processes including synthesis, release, reuptake, degradation, and receptor activities (Beretta et al., 2025; Newell & Patisaul, 2025). The developmental elevation in serotonin during larval stages may disrupt the normal trajectories of serotonergic circuit formation and synaptic plasticity (Winberg & Thörnqvist, 2016). The pronounced upregulation of serotonin receptors and transporters following BPS exposure suggests compensatory or maladaptive responses to elevated intrasynaptic serotonin concentrations. In the normal zebrafish brain, serotonin receptors are distributed throughout multiple brain regions including the hypothalamus, preoptic area, and dorsal pallium, regions critical for social behavior and emotional regulation (Corradi & Filosa, 2021; Gabriel et al., 2009; Montgomery et al., 2018). The upregulation of postsynaptic serotonin receptors (*htr1aa, htr1ab, htr1b, htr1d, htr2aa, htr2b*) may reflect a compensatory mechanism attempting to maintain homeostatic serotonergic signaling in the face of elevated 5-HT. Alternatively, receptor upregulation could indicate desensitization or downregulation of receptor responsiveness, a phenomenon commonly observed when neurons are chronically exposed to elevated neurotransmitter levels (Allen et al., 2008; Raote et al., 2007). The dramatic upregulation of serotonin transporters (*slc6a4a and slc6a4b*) is particularly noteworthy, as these proteins are responsible for the reuptake of 5-HT from the synaptic cleft back into presynaptic neurons (Huang et al., 2023; Yuan et al., 2014). The 8-fold increase in *slc6a4b* expression suggests an attempt by the CNS to clear excess serotonin and restore homeostatic balance. However, this compensatory upregulation may be insufficient to normalize serotonergic signaling, resulting in persistently elevated 5-HT levels and dysregulated neurotransmission. Molecular docking of BPS against six serotonergic and endocrine targets provided additional mechanistic support: the highest predicted binding affinity was observed for the HTR1A receptor (-6.274 kcal/mol; PDB: 7E2Y), followed by TPH2 (-5.805 kcal/mol) and ERα (-5.778 kcal/mol), indicating that BPS can directly engage both serotonin receptors and the rate-limiting serotonin synthesis enzyme. Furthermore, SLC6A4 (SERT) ranked fourth in network hub score (0.389), consistent with the 8-fold upregulation of *slc6a4b* observed experimentally and supporting direct BPS-SERT interaction as a plausible mechanism underlying elevated synaptic serotonin. Monoamine GPCR pathways (WikiPathways; FDR = 1.47 × 10^-^⁷) were also among the top enriched terms in the expanded 61-gene CTD-derived novel target network, reinforcing the serotonergic synapse as the primary pharmacological target of BPS.

This mechanistic framework is strongly supported by the observed impairment in social behaviour, where BPS-exposed larvae showed a significant reduction in time spent in the conspecific zone. Social behaviour in zebrafish is tightly regulated by serotonergic signaling within brain regions such as the dorsal pallium and hypothalamus (Herculano & Maximino, 2014). Dysregulation of this system, even in the absence of overt oxidative damage or widespread apoptosis, can lead to selective behavioural deficits. Therefore, the present results support the conclusion that BPS primarily exerts its neurodevelopmental toxicity through serotonergic dysregulation, with oxidative stress acting as a contributing downstream damage pathway. The experimental and computational findings converge on a unified mechanistic model: BPS directly binds serotonin receptors and TPH2 with measurable affinity, elevates synaptic 5-HT, and dysregulates BDNF-CREB1 hub signalling, culminating in serotonergic circuit immaturity and social behavioural deficits. The integration of PPI network topology, pathway enrichment, and molecular docking provides convergent, multi-scale evidence supporting the mechanistic framework of this study, and identifies HTR1A, TPH2, BDNF, and CREB1 as key regulatory targets underlying BPS-induced neurodevelopmental and behavioural disruption.

## 5. Conclusion

The present study demonstrates that developmental exposure to an environmentally relevant concentration of BPS (30 µg/L) induces significant social behavioural deficits in zebrafish larvae, driven primarily by serotonergic dysregulation. Despite the absence of increased lipid peroxidation, BPS exposure triggered a disrupted antioxidant response, suppression of key apoptotic regulators, and robust upregulation of serotonergic receptors and transporters, indicating functional reprogramming of neurodevelopmental pathways. Integration of experimental and computational analyses revealed that BPS-induced molecular changes are organized within a highly interconnected network centered on BDNF and CREB1, linking serotonergic signaling, oxidative stress responses, and neurotrophic regulation. Pathway enrichment consistently identified the serotonergic synapse as the most affected biological pathway, while molecular docking provided direct evidence that BPS can bind multiple serotonin receptors and synthesis-related proteins, offering a mechanistic basis for altered neurotransmission. These findings support a model in which BPS acts as a multi-target neuroactive compound, perturbing neurotransmitter systems and regulatory networks at multiple levels. This work advances our understanding of BPS neurotoxicity by establishing a systems-level framework connecting molecular, biochemical, and behavioural effects, and identifies key molecular targets (HTR1A, TPH2, BDNF, CREB1) for future investigations. Given the widespread environmental presence of BPS, these results raise important concerns regarding its potential ecological and neurodevelopmental impacts.

## CRediT authorship contribution statement

**A K M Munzurul Hasan:** Conceptualization, Methodology, Software, Formal analysis, Investigation, Data Curation, Writing-original draft preparation, Writing-review and editing, Visualization, Validation; **Mahesh Rachamalla:** Investigation; **Som Niyogi:** Writing-review and editing, Resources; Supervision, Project administration, Funding acquisition; **Douglas P. Chivers:** Writing-review and editing, Resources; Supervision, Project administration, Funding acquisition.

## Acknowledgments

A K M Munzurul Hasan gratefully acknowledges the financial support received through the University Graduate Scholarship (UGS) and Graduate Teaching Fellowship (GTF) programs at the University of Saskatchewan. This work was funded by Discovery Grants from the Natural Sciences and Engineering Research Council of Canada (NSERC) awarded to D.P.C. and S.N. Imaging analyses were performed using instrumentation provided by the Microscopy Core Facility, Department of Biology, University of Saskatchewan.

## Reference

Allen, J. A., Yadav, P. N., & Roth, B. L. (2008). Insights into the regulation of 5-HT2A serotonin receptors by scaffolding proteins and kinases. Neuropharmacology, 55(6), 961–968. 10.1016/j.neuropharm.2008.06.048

Anwar, S., Alrumaihi, F., Sarwar, T., Babiker, A. Y., Khan, A. A., Prabhu, S. V., & Rahmani, A. H. (2024). Exploring therapeutic potential of catalase: Strategies in disease prevention and management. Biomolecules, 14(6), 697.

Attaran, A., Salahinejad, A., Naderi, M., Crane, A. L., Chivers, D. P., & Niyogi, S. (2021). Transgenerational effects of selenomethionine on behaviour, social cognition, and the expression of genes in the serotonergic pathway in zebrafish. Environmental Pollution, 286, 117289. 10.1016/j.envpol.2021.117289

Bacqué-Cazenave, J., Bharatiya, R., Barrière, G., Delbecque, J.-P., Bouguiyoud, N., Di Giovanni, G., Cattaert, D., & De Deurwaerdère, P. (2020). Serotonin in animal cognition and behavior. International Journal of Molecular Sciences, 21(5), 1649.

Benito, E., & Barco, A. (2015). The neuronal activity-driven transcriptome. Molecular Neurobiology, 51(3), 1071–1088.

Beretta, E., Cuboni, G., & Deidda, G. (2025). Unveiling GABA and serotonin interactions during neurodevelopment to re-open adult critical periods for neuropsychiatric disorders. International Journal of Molecular Sciences, 26(12), 5508.

Carlezon, W. A., Duman, R. S., & Nestler, E. J. (2005). The many faces of CREB. Trends in neurosciences, 28(8), 436–445.

Corradi, L., & Filosa, A. (2021). Neuromodulation and behavioral flexibility in larval zebrafish: from neurotransmitters to circuits. Frontiers in molecular neuroscience, 14, 718951.

Cunha, C., Brambilla, R., & Thomas, K. L. (2010). A simple role for BDNF in learning and memory? Frontiers in molecular neuroscience, 3, 865.

Dworkin, S., Heath, J. K., deJong-Curtain, T. A., Hogan, B. M., Lieschke, G. J., Malaterre, J., Ramsay, R. G., & Mantamadiotis, T. (2007). CREB activity modulates neural cell proliferation, midbrain–hindbrain organization and patterning in zebrafish. Developmental biology, 307(1), 127–141.

Elmore, S. (2007). Apoptosis: a review of programmed cell death. Toxicologic pathology, 35(4), 495–516.

Fabrello, J., & Matozzo, V. (2022). Bisphenol analogs in aquatic environments and their effects on marine species—a review. Journal of Marine Science and Engineering, 10(9), 1271.

Firdous, S. M., Pal, S., Khanam, S., & Zakir, F. (2024). Behavioral neuroscience in zebrafish: Unravelling the complexity of brain-behavior relationships. Naunyn-Schmiedeberg’s Archives of Pharmacology, 397(12), 9295–9313.

Gabriel, J. P., Mahmood, R., Kyriakatos, A., Söll, I., Hauptmann, G., Calabrese, R. L., & El Manira, A. (2009). Serotonergic modulation of locomotion in zebrafish—endogenous release and synaptic mechanisms. Journal of Neuroscience, 29(33), 10387–10395.

Gupta, S., & Haldar, C. (2017). Short day length enhances physiological resilience of the immune system against 2-deoxy-d-glucose-induced metabolic stress in a tropical seasonal breeder Funambulus pennanti. Hormones and Behavior, 89, 157–166. 10.1016/j.yhbeh.2017.01.004

Hasan, A. K. M. M., Rachamalla, M., Uddin, M. H., Putnala, S. K., Krol, E. S., Niyogi, S., & Chivers, D. P. (2026). Developmental exposure to Bisphenol S causes neurobehavioural deficits in larval zebrafish (Danio rerio). Aquatic Toxicology, 291, 107701. 10.1016/j.aquatox.2025.107701

Herculano, A. M., & Maximino, C. (2014). Serotonergic modulation of zebrafish behavior: Towards a paradox. Progress in Neuro-Psychopharmacology and Biological Psychiatry, 55, 50–66. 10.1016/j.pnpbp.2014.03.008

Huang, C., van Wijnen, A. J., & Im Sampen, H.-J. (2023). Serotonin transporter (5-HTT, SERT, SLC6A4) and sodium-dependent reuptake inhibitors (SRIs) as modulators of pain behaviors and analgesic responses. The Journal of Pain, 25(3), 618.

Ighodaro, O. M., & Akinloye, O. A. (2018). First line defence antioxidants-superoxide dismutase (SOD), catalase (CAT) and glutathione peroxidase (GPX): Their fundamental role in the entire antioxidant defence grid. Alexandria Journal of Medicine, 54(4), 287–293. 10.1016/j.ajme.2017.09.001

Jayanthi, L. D., & Ramamoorthy, S. (2025). Role of Phosphorylation of Serotonin and Norepinephrine Transporters in Animal Behavior: Relevance to Neuropsychiatric Disorders. International Journal of Molecular Sciences, 26(16), 7713.

Ji, K., Hong, S., Kho, Y., & Choi, K. (2013). Effects of bisphenol S exposure on endocrine functions and reproduction of zebrafish. Environmental Science & Technology, 47(15), 8793–8800.

Ji, L. L. (2002). Exercise-induced modulation of antioxidant defense. Annals of the New York Academy of Sciences, 959(1), 82–92.

Jomova, K., Alomar, S. Y., Alwasel, S. H., Nepovimova, E., Kuca, K., & Valko, M. (2024). Several lines of antioxidant defense against oxidative stress: antioxidant enzymes, nanomaterials with multiple enzyme-mimicking activities, and low-molecular-weight antioxidants. Archives of Toxicology, 98(5), 1323–1367.

Kamat, P. K., Kalani, A., Rai, S., Swarnkar, S., Tota, S., Nath, C., & Tyagi, N. (2016). Mechanism of oxidative stress and synapse dysfunction in the pathogenesis of Alzheimer’s disease: understanding the therapeutics strategies. Molecular Neurobiology, 53(1), 648–661.

Lee, B., Cao, R., Choi, Y. S., Cho, H. Y., Rhee, A. D., Hah, C. K., Hoyt, K. R., & Obrietan, K. (2009). The CREB/CRE transcriptional pathway: protection against oxidative stress-mediated neuronal cell death. Journal of neurochemistry, 108(5), 1251–1265.

Lee, S., Suk, K., Kim, I. K., Jang, I. S., Park, J. W., Johnson, V. J., Kwon, T. K., Choi, B. J., & Kim, S. H. (2008). Signaling pathways of bisphenol A–induced apoptosis in hippocampal neuronal cells: Role of calcium-induced reactive oxygen species, mitogen-activated protein kinases, and nuclear factor–κB. Journal of neuroscience research, 86(13), 2932–2942.

Li, B., Huang, Y., Pi, D., Li, X., Guo, Y., Liang, Z., Song, X., Wang, J., & Wang, X. (2024). Effects of acute and developmental exposure to bisphenol S on Chinese medaka (Oryzias sinensis). Journal of Xenobiotics, 14(2), 452–466.

Liao, C., Liu, F., Alomirah, H., Loi, V. D., Mohd, M. A., Moon, H.-B., Nakata, H., & Kannan, K. (2012). Bisphenol S in urine from the United States and seven Asian countries: occurrence and human exposures. Environmental Science & Technology, 46(12), 6860–6866.

Lu, B., Nagappan, G., Guan, X., Nathan, P. J., & Wren, P. (2013). BDNF-based synaptic repair as a disease-modifying strategy for neurodegenerative diseases. Nature Reviews Neuroscience, 14(6), 401–416.

Maćczak, A., Cyrkler, M., Bukowska, B., & Michałowicz, J. (2017). Bisphenol A, bisphenol S, bisphenol F and bisphenol AF induce different oxidative stress and damage in human red blood cells (in vitro study). Toxicology in Vitro, 41, 143–149.

Manzoor, M. F., Tariq, T., Fatima, B., Sahar, A., Tariq, F., Munir, S., Khan, S., Nawaz Ranjha, M. M. A., Sameen, A., & Zeng, X.-A. (2022). An insight into bisphenol A, food exposure and its adverse effects on health: A review. Frontiers in nutrition, 9, 1047827.

Miller, N., & Gerlai, R. (2012). From schooling to shoaling: patterns of collective motion in zebrafish (Danio rerio). Plos One, 7(11), e48865.

Molina-Molina, J.-M., Amaya, E., Grimaldi, M., Sáenz, J.-M., Real, M., Fernández, M. F., Balaguer, P., & Olea, N. (2013). In vitro study on the agonistic and antagonistic activities of bisphenol-S and other bisphenol-A congeners and derivatives via nuclear receptors. Toxicology and Applied Pharmacology, 272(1), 127–136. 10.1016/j.taap.2013.05.015

Montgomery, J. E., Wahlstrom-Helgren, S., Wiggin, T. D., Corwin, B. M., Lillesaar, C., & Masino, M. A. (2018). Intraspinal serotonergic signaling suppresses locomotor activity in larval zebrafish. Developmental Neurobiology, 78(8), 807–827.

Moreman, J., Lee, O., Trznadel, M., David, A., Kudoh, T., & Tyler, C. R. (2017). Acute toxicity, teratogenic, and estrogenic effects of bisphenol A and its alternative replacements bisphenol S, bisphenol F, and bisphenol AF in zebrafish embryo-larvae. Environmental Science & Technology, 51(21), 12796–12805.

Narasimhamurthy, R. K., Andrade, D., & Mumbrekar, K. D. (2022). Modulation of CREB and its associated upstream signaling pathways in pesticide-induced neurotoxicity. Molecular and Cellular Biochemistry, 477(11), 2581–2593. 10.1007/s11010-022-04472-7

Newell, A. J., & Patisaul, H. B. (2025). Modeling the developing nervous system: a neuroscience perspective on the use of new approach methodologies in developmental neurotoxicity testing. Toxicological Sciences, 205(2), 245–273.

Qiu, W., Zhao, Y., Yang, M., Farajzadeh, M., Pan, C., & Wayne, N. L. (2016). Actions of bisphenol A and bisphenol S on the reproductive neuroendocrine system during early development in zebrafish. Endocrinology, 157(2), 636–647.

Raote, I., Bhattacharya, A., & Panicker, M. M. (2007). Serotonin 2A (5-HT2A) receptor function: ligand-dependent mechanisms and pathways. Serotonin receptors in neurobiology.

Rosenmai, A. K., Dybdahl, M., Pedersen, M., Alice van Vugt-Lussenburg, B. M., Wedebye, E. B., Taxvig, C., & Vinggaard, A. M. (2014). Are Structural Analogues to Bisphenol A Safe Alternatives? Toxicological Sciences, 139(1), 35–47. 10.1093/toxsci/kfu030

Saili, K. S., Corvi, M. M., Weber, D. N., Patel, A. U., Das, S. R., Przybyla, J., Anderson, K. A., & Tanguay, R. L. (2012). Neurodevelopmental low-dose bisphenol A exposure leads to early life-stage hyperactivity and learning deficits in adult zebrafish. Toxicology, 291(1-3), 83–92.

Salahinejad, A., Attaran, A., Naderi, M., Meuthen, D., Niyogi, S., & Chivers, D. P. (2021). Chronic exposure to bisphenol S induces oxidative stress, abnormal anxiety, and fear responses in adult zebrafish (Danio rerio). Science of The Total Environment, 750, 141633.

Salahinejad, A., Naderi, M., Attaran, A., Meuthen, D., Niyogi, S., & Chivers, D. P. (2020). Effects of chronic exposure to bisphenol-S on social behaviors in adult zebrafish: Disruption of the neuropeptide signaling pathways in the brain. Environmental Pollution, 262, 113992. 10.1016/j.envpol.2020.113992

Semeão, K. A., Dutra-Tavares, A. C., Ribeiro-Carvalho, A., Isnardo-Fernandes, J., Lopes, L. D., Souza, G. S., Nunes-Freitas, A. L., Silva, B. S., Filgueiras, C. C., & Manhães, A. C. (2025). Serotonergic and Cholinergic Imbalance in the Offspring of Rats Exposed to Bisphenol A and Bisphenol S During Pregnancy and Lactation: Short-and Long-Term Effects. International Journal of Molecular Sciences, 26(19), 9329.

Shehna Mahim, S., Anjali, V., Remya, V., Reshmi, S., & Aruna Devi, C. (2021). Oxidative stress responses of a freshwater fish, Labeo rohita, to a xenobiotic, bisphenol S. Journal of Biochemical and Molecular Toxicology, 35(8), e22820.

Wan, Y., Xia, W., Yang, S., Pan, X., He, Z., & Kannan, K. (2018). Spatial distribution of bisphenol S in surface water and human serum from Yangtze River watershed, China: Implications for exposure through drinking water. Chemosphere, 199, 595–602. 10.1016/j.chemosphere.2018.02.040

Wang, W., Li, Z., Zhang, X., Zhang, J., & Ru, S. (2022). Bisphenol S impairs behaviors through disturbing endoplasmic reticulum function and reducing lipid levels in the brain of zebrafish. Environmental Science & Technology, 57(1), 582–594.

Weber, D. N., Hoffmann, R. G., Hoke, E. S., & Tanguay, R. L. (2015). Bisphenol A exposure during early development induces sex-specific changes in adult zebrafish social interactions. *Journal of Toxicology and Environmental Health*, Part A, 78(1), 50–66.

Whitaker-Azmitia, P. M. (2001). Serotonin and brain development: role in human developmental diseases. Brain research bulletin, 56(5), 479–485.

Winberg, S., & Thörnqvist, P.-O. (2016). Role of brain serotonin in modulating fish behavior. Current Zoology, 62(3), 317–323. 10.1093/cz/zow037

Yamazaki, E., Yamashita, N., Taniyasu, S., Lam, J., Lam, P. K. S., Moon, H.-B., Jeong, Y., Kannan, P., Achyuthan, H., Munuswamy, N., & Kannan, K. (2015). Bisphenol A and other bisphenol analogues including BPS and BPF in surface water samples from Japan, China, Korea and India. Ecotoxicology and Environmental Safety, 122, 565–572. 10.1016/j.ecoenv.2015.09.029

Yang, Y., Yang, Y., Zhang, J., Shao, B., & Yin, J. (2019). Assessment of bisphenol A alternatives in paper products from the Chinese market and their dermal exposure in the general population. Environmental Pollution, 244, 238–246.

Yuan, J., Kang, C., Wang, M., Wang, Q., Li, P., Liu, H., Hou, Y., Su, P., Yang, F., & Wei, Y. (2014). Association study of serotonin transporter SLC6A4 gene with Chinese Han irritable bowel syndrome. Plos One, 9(1), e84414.

